# Cellular Senescence Drives Zinc Accumulation and Transporter Dysregulation in Intestinal Epithelial Cells

**DOI:** 10.64898/2026.02.19.706872

**Authors:** Kristofer C Terrell, Suyun Choi, Jiahn Choi, Sangyong Choi

**Affiliations:** Department of Nutritional Sciences, University of Connecticut, Storrs, CT, USA; Department of Medicine and Department of Molecular Pharmacology, Albert Einstein College of Medicine, Bronx, NY, USA

**Keywords:** aging, zinc, cellular senescence, intestine, nutrition, Golgi, gene regulation

## Abstract

Zinc is essential for life, and its regulation is tightly controlled by numerous transporters. As we age, our micronutrient levels, intake, and absorption change. Additionally, senescent cells increase with age and can contribute to the progression of age-related diseases. The study of Zn homeostasis in senescent intestinal cells is a relatively unexplored area that we aimed to investigate. Using two models to induce senescence in intestinal epithelial cells—etoposide treatment and γ-irradiation—we observed that Zn levels increased in the cells, likely due to the upregulation of Zn transporters ZIP4 and ZnT7. This upregulated Zn seems to accumulate in the Golgi apparatus, and when Zn accumulation is blocked through chelation, a rescue effect occurs, marked by a decrease in senescence markers. This research emphasizes the role of Zn in senescent cells and its possible involvement in the development of senescence and the disrupted Zn homeostasis seen with aging.

## Introduction

As we age, various molecular, cellular, and systemic processes are altered, leading to a loss of normal function and eventually death. Various hallmarks are associated with aging, including genomic instability, mitochondrial dysfunction, cellular senescence, stem cell exhaustion, and chronic inflammation or inflammaging [1]. Cellular senescence, the stable arrest of the cell cycle, has been seen to increase with aging, progress many age-related diseases, and alter the gastrointestinal tract [2-4]. The functionality of the gastrointestinal system affects nearly all parts of the human body, absorbing key nutrients required for life, functioning as a barrier, and housing a microbiome that contains over 150 times more genetic information than the human genome, and is thought to be the most significant microbiome across the body for maintaining health.

Zinc (Zn) is an essential mineral that catalyzes numerous reactions. Over 3,000 human proteins, or ≥ 10% of all human proteins, employ Zn as a cofactor [5]. To maintain Zn levels in the body, Zn must be consumed in the diet and absorbed across the intestinal epithelium. If consumption is too low or absorption is dysregulated, the risk of Zn deficiency increases. An estimated 17.3% of the world’s population is at risk of inadequate Zn intake [6]. Within the at-risk population, young children and elderly adults are most at risk, alongside pregnant and lactating women. Deficiency in Zn is linked with altered immune function, growth retardation, and various other outcomes due to its wide-reaching function mentioned previously [7-9].

Screening studies for Zn status, measured via serum Zn levels, in populations above 60, have found the prevalence of deficiency to be approximately 8.68-10.1% or 90 million people worldwide [10,11]. Some of this can be attributed to dietary pattern shifts. However, dietary absorption may also play a role in the prevalence of Zn deficiency. Early studies, predominantly in men, found that there was decreased absorption of Zn measured via stable isotope feedings in the same age ranges as the previously mentioned studies [12,13]. This highlights that there may be a change in Zn transport at the epithelium level.

There are several changes seen in the GI system as we age [14]. In model organisms, there are changes in villi height, crypt depth, microbiome composition, and importantly, changes in macronutrient absorption [15]. Moreover, some of these changes can be rescued upon removal of senescent cells [4,16].

Zn accumulates as a function of age in tissues such as liver and muscle, but decreases in serum, kidney, and bladder *in vivo*[17,18]. Like Zn, various other trace metals are known to accumulate in tissues with aging and have been found to be dysregulated in senescent cells [19-21]. However, the role of Zn in senescent cells is not fully understood.

### Primary objective

The primary objective of this study was to determine the effect of cellular senescence on Zn homeostasis in a gut epithelial cell model and how this Zn regulation may contribute to aging-related Zn deficiency. Secondary objectives were to elucidate the specific role that Zn may play within the development of senescence.

## Methods

### Cell Culture and Treatment

C2BBe1 (clone of Caco-2) brush border-expressing enterocyte cells were kindly gifted from Dr. Zui Pan (University of Texas at Arlington). C2BBe1 enterocytes were grown in Dulbecco’s modified Eagle Medium (Corning: 10-013-CV) containing 10% fetal bovine serum (Sigma-Aldrich: F1051) and 1% penicillin/streptomycin (Corning: 30002CI) in a humidified incubator with 5% CO2 at 37°C. To induce senescence via etoposide, we used a previously reported method with slight modifications [22]. 24 hours post seeding, cells were treated with media containing 5μM etoposide or an equal volume of dimethyl sulfoxide for 48 hours, then replaced back to regular media, and allowed to recover for 48 hours until collected, splitting the dimethyl sulfoxide group at 24 hours post recovery to prevent excessive cell confluence. To induce senescence using irradiation, C2BBe1 cells were seeded, then 24 hours later were exposed to gamma irradiation at 5 gray (Cesium-137). 48 hours later, cells were collected for analysis.

### Cell Viability Assay

Cell viability was measured for both methods of senescence induction, etoposide or irradiation, determined by the MTT (3-(4,5-dimethylthiazol-2-yl)-2,5-diphenyl tetrazolium bromide) assay as described previously [23].For etoposide treatments, cells were seeded onto a 96-well plate and treated with various concentrations of etoposide. For the irradiation experiment, cells were seeded onto 35 mm dishes and exposed to various irradiation intensities. For both, multiple time points were collected. Media containing 2.4 mM MTT was added to the cells for 3 hours. The media was removed, and 150 μL of DMSO was added to each well. Absorbance was read at 570 nm.

### Senescence-Associated β-Galactosidase Staining and Quantification

Senescence-associated β-Galactosidase staining was performed as described by Debacq-Chainiaux *et al*. [24]. Briefly, cells were washed twice with phosphate-buffered saline, fixed in paraformaldehyde for 15 minutes at room temperature, and incubated overnight at 37°C in staining solution. The activity of the staining solution was imaged in brightfield using a Lionheart^™^ FX imager. Quantification of positively stained cells was done as described by Hooten and Evans [25]. The number of positive cells within total cells (>100) was counted to calculate the percentage of positively stained cells for each of the three biological replicates.

### Cellular Zn Quantification

Two methods of Zn quantification were used. First, Zn was measured using the fluorescent probe Zinpyr-1 (Cayman Chemical 15122). Cells went through both methods of senescence induction and were then stained with 5μM Zinpyr-1 for 30 minutes, and imaged using a Lionheart^™^ FX imager. Cellular analysis was done in the Gen5 software measuring mean fluorescence intensity within individual cells on the raw, unaltered image. Images presented in this paper have a background reduction step applied, also using Gen5 software. The second method of Zn assessment was using a colorimetric assay. Cellular Zn levels were quantified using a colorimetric Zn assay kit (Sigma-Aldrich: MAK032), normalized to total protein.

### Fluorescence Staining

Cells grown on glass chamber slides were tri-stained with Zinpyr-1, BODIPY^™^ TR Ceramide (ThermoFisher: D7540), and mounted with SlowFade^™^ Glass Soft-set Antifade Mountant, with DAPI (ThermoFisher: S36920). Cells are first stained with Zinpyr-1 as previously described, then washed and stained for 30 minutes with 5 μM ceramide, then washed and mounted with DAPI-containing mountant. These were then used for 60x oil-immersion microscopy using the Lionheart^™^ FX imager.

### Gene expression analysis: RNA Extraction, cDNA Synthesis, and RT-qPCR

Total RNA was extracted using Aurum^™^ Total RNA Mini Kit (BioRad: 7326820). Reverse transcription was done using iScript cDNA Synthesis Kit (BioRad: 1708890). Real time quantitative PCR (RT-qPCR) was preformed with a CFX Opus 96 Real-Time PCR system using iTaq Universal SYBR Green supermix (BioRad: 1725124). Expression relative to housekeeping gene GAPDH was used for all experiments. Primers used for RT-qPCR can be found in Supplementary Table 1.

A single-cell RNA sequencing dataset was analyzed to investigate Zn transporter genes across various cell lineages in the small intestine epithelium, as previously described [26]. Briefly, CB6F1 hybrid male mice were harvested at either 5 or 24 months of age following a three-month gerotherapeutic intervention with rapamycin or metformin. Intestines were collected, and single-cell RNA sequencing libraries were constructed using the 3’ kit protocols from 10X Genomics (San Francisco, CA). Sequences were aligned and converted to count matrices, and quality control and downstream analysis were performed in R. Differential gene expression among experimental groups was analyzed using the Seurat FindMarkers function.

### Western Blotting

Total protein was extracted using Membrane and cytosolic fractions were extracted using RIPA lysis and Extraction Buffer (ThermoFisher Scientific 89900). 50 μg protein was loaded onto handmade 10-12% SDS gel and transferred onto 0.2 μm polyvinylidene fluoride membrane. Membranes were blocked 1hr room temp in 10% non-fat dairy milk, rinsed, and incubated rocking overnight with the primary antibody at 4°C. Secondary antibodies were incubated one hour at room temperature then washed and excited via Pierce^™^ ECL Western Blotting Substrate (ThermoFisher 32106), then imaged and quantified using Bio Rad’s ChemiDoc and Imagelab software. Primary antibodies included, Beta-Actin (Proteintech 60008-1-lg),Lamin B1 (Proteintech 66095-1-lg), p21 (Proteintech 10355-1-AP), Zip4 (Bioss bs-19832R), ZnT7 (Proteintech 13966-1-AP).

### FRET Microscopy

pcDNA-Golgi-ZapCY1 and pcDNA-ER-ZAPCY1 plasmids were a gift from Dr. Amy Palmer (Addgene plasmid # 36322 and #36319). Fluorescence Resonance Energy Transfer (FRET) imaging was done using the Lionheart^™^ FX imager. Cells were transfected with either the Golgi apparatus and the ER-specific Zn sensor, respectively [27]. These cells were then treated with etoposide as described previously, and imaged every 20 seconds, with a treatment of 20 μM ZnSO_4_:pyrithione 1:2, where pyrithione is 2-meracaptopyridine-N-oxide, followed after by 100uM treatment of a Zn-specific chelator, tris(2-pyridylmethyl)amine (TPA) [28].

### Zn Depletion

Zn chelation was performed using the Zn specific membrane permeable tris(2-pyridylmethyl)amine (TPA). TPA was treated to cells 24 hrs after seeding, and refreshed with each media swap, following the etoposide administration methods mentioned above.

### Statistical Analysis

All data are expressed as means ± standard deviation. The Student’s *t*-test or one-way analysis of variance (ANOVA), or two-way ANOVA, were used to evaluated differences between groups. A value of P < 0.05 was considered significantly significant. Statistical analysis was performed using GraphPad Prism software version 10.4.2.

## Results

### Inducing senescence in C2BBe1 Cells

To examine the effect of senescence on the gut epithelium, C2BBe1 cells were treated with etoposide or gray irradiation, as a method to induce DNA replication inhibition or direct DNA damage. Various treatment dosages and recovery times were examined to optimize the condition of inducing senescence using cell viability (Supplemental Fig. 1) and further analyses. As the cells exhibited a lack of proliferation without undergoing cell death, the viability assay results led us to employ 5 μM etoposide or 5 Gy irradiation, with sample collection occurring at the 48 h recovery time. Senescence-associated β-galactosidase staining showed a significant increase in the staining-positive cells in both etoposide and irradiated cells (Fig. 1A-B). Protein expression of senescence-associated proteins further supports increased senescence induction, marked by decreases of Lamin B1 and increased p21 (Fig. 1C-F) in both cell-senescence models. Expression of various senescence-associated genes was also altered following typical senescence hallmarks. Increased CDKN1A, IL6, TIMP1, and a decrease of LMNB1 were seen in both methods of senescence induction (Fig. 1 G-H). With senescence induction validated, we explored Zn homeostasis within these cells.

**Fig. 1.**
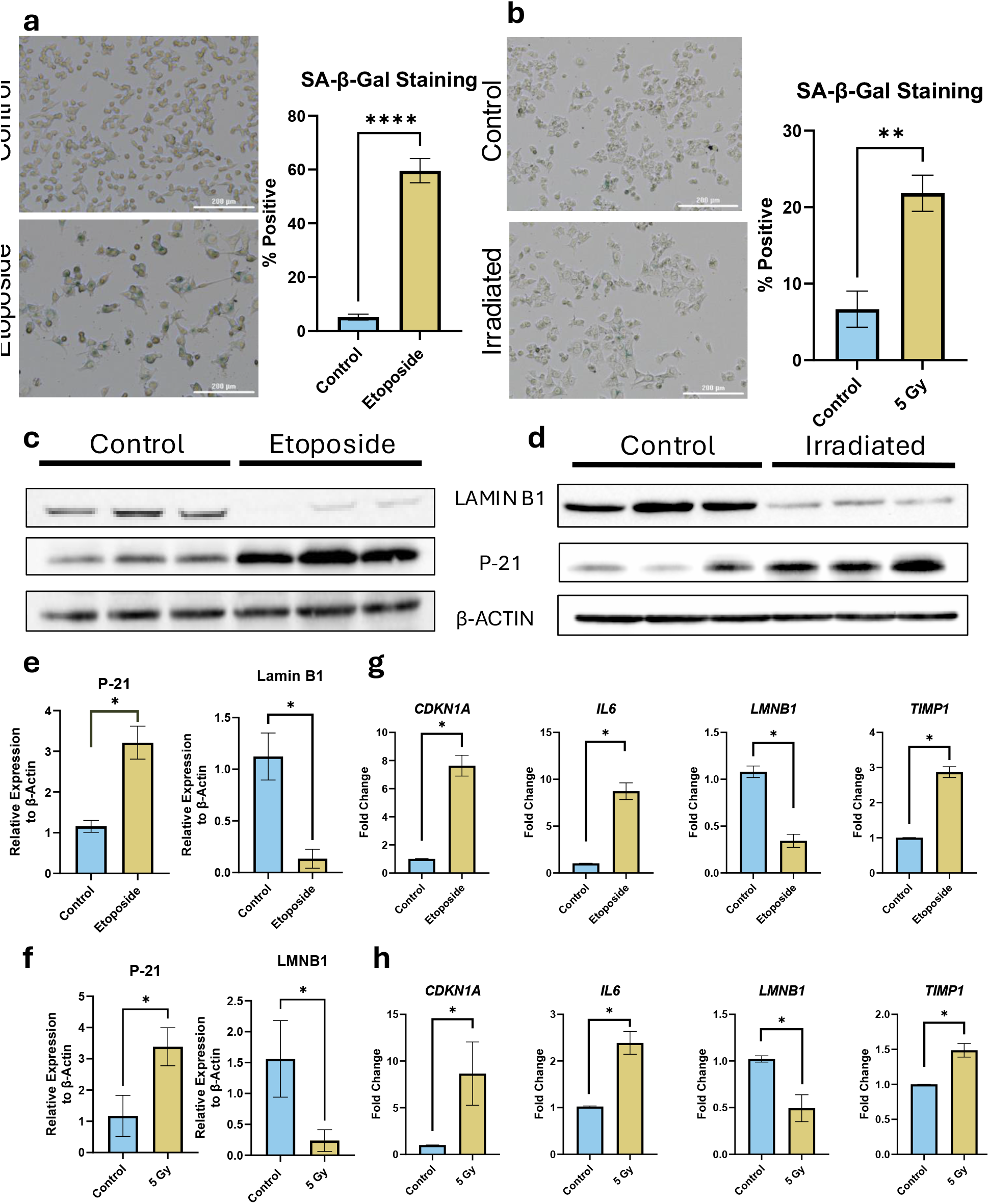
Senescence induction validation for etoposide or γ-irradiation. Representative images of senescence-associated ϐ-galactosidase staining at 48 hours recovery between control (top) or treated cells (bottom) and their quantification for etoposide (A) or irradiation-induced senescence (B). Scale bar = 200 μm. Western blot analysis of p21 and LaminB1, using ϐ-actin as a loading control in etoposide-treated (C) or irradiated cells (D), along with their corresponding quantification (E-F). RNA expression of senescent-associated genes *CDKN1A, IL6, LMNB1*, and *TIMP1* was measured by quantitative RT-qPCR and normalized with housekeeping gene *GAPDH* for etoposide (G) or irradiation (H). Data represents mean values ± standard deviation from at least n=3 independent experiments are depicted. For all analyses, a p < 0.05 was considered significant.

### Zn homeostasis in senescent and aged enterocytes

To examine Zn status in senescent C2BBE1 cells, we quantified liable Zn using Zinpyr-1, a fluorescent Zn dye, and total cellular Zn using a colorimetric kit. Both labile Zn and total Zn increased upon senescence inductions (Fig. 2A-D). To understand how this Zn was entering the cells and the regulation of Zn transporters, we performed RT-qPCR on highly expressed Zn transporters, displayed as a heatmap for both treatments (Fig. 3A-B). The genes that had significant changes in one or both of the senescence induction methods are shown to the right of the heatmap. In both etoposide-treated and irradiated cells, *SLC39A4* and *SLC30A7* were significantly upregulated. *SLC39A4, SLC39A11, SLC39A13, SLC30A1*, and *SLC30A4* were increased in etoposide-treated cells; however, in irradiated cells, these genes did not reach significance. *SLC30A6* was also slightly decreased in etoposide-treated cells. However, this was not matched in irradiated cells. *SLC39A14* increased in irradiated cells, but did not reach significance in etoposide-treated cells. Following trends seen in gene expression, we observed that protein expression of ZIP4 and ZnT7 was also increased in etoposide-induced senescence and irradiation-induced senescence (Fig. 4A-B). Similarly, in aged mice compared to young mice, *Slc39a4* expression increases in multiple enterocyte lineage cells—EC1, 2, 3, and 6— (supplementary fig. 2 a-c) in the single-cell RNA-seq dataset. Furthermore, treatment with gerotherapeutics, such as rapamycin or metformin, has the opposite effect, reducing *Slc39a4* expression.

**Fig. 2.**
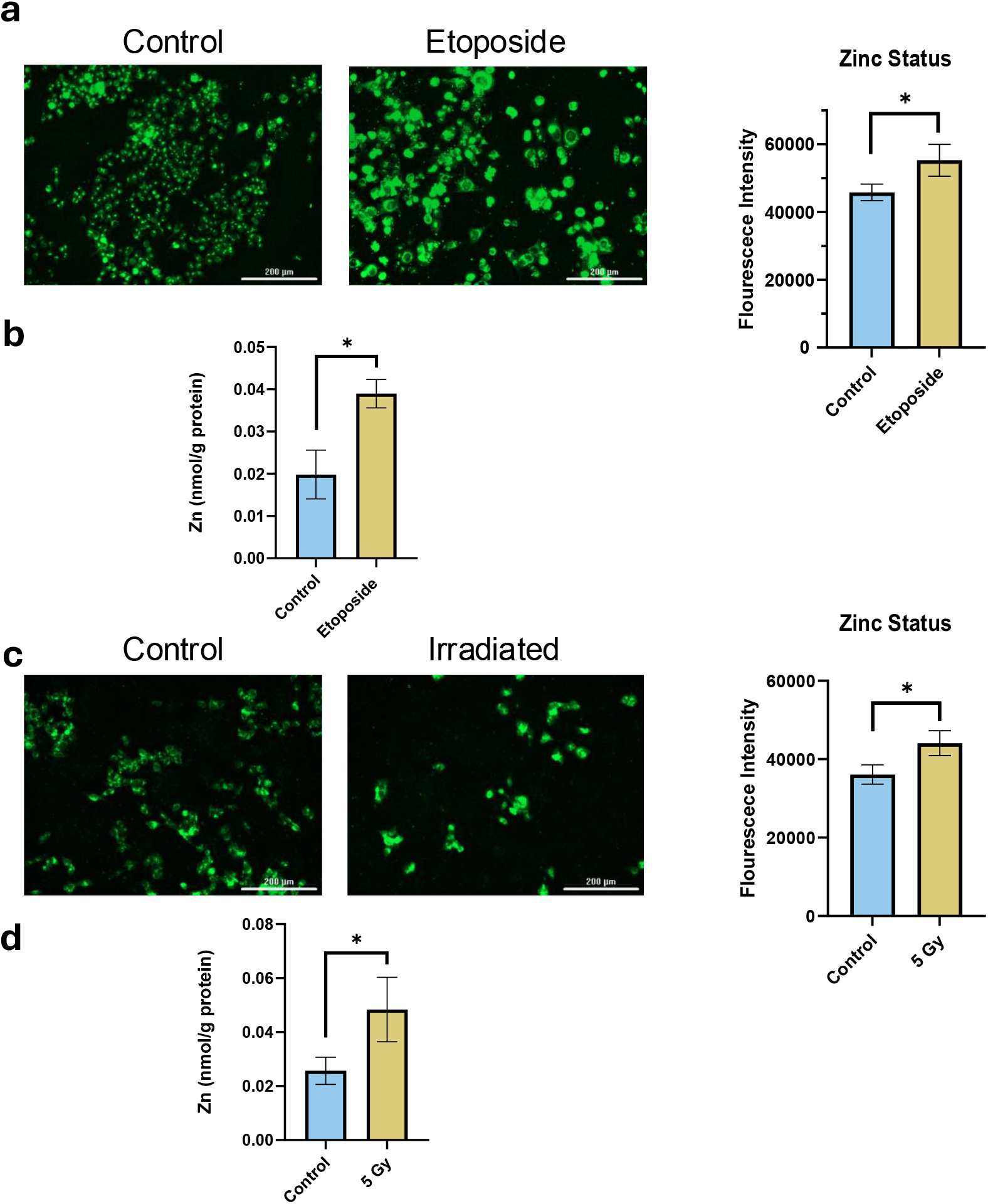
Zn status in senescent cells. Representative images (left) and their quantification (right) fluorescence intensity of Zinpyr-1, a Zn-specific dye, between control and etoposide-induced senescence (A). Quantitative colorimetric Zn assay in etoposide-induced senescence (B). Representative images (left) and their quantification (right) of fluorescence intensity of Zinpyr-1, in irradiation-induced senescence (C). Quantitative colorimetric Zn assay in irradiation-induced senescence (D). For fluorescent quantification, n ≥ 40 cells. Scale bars = 200 μm. Data represents mean ± standard deviation. For analyses, a p < 0.05 was considered significant.

**Fig. 3.**
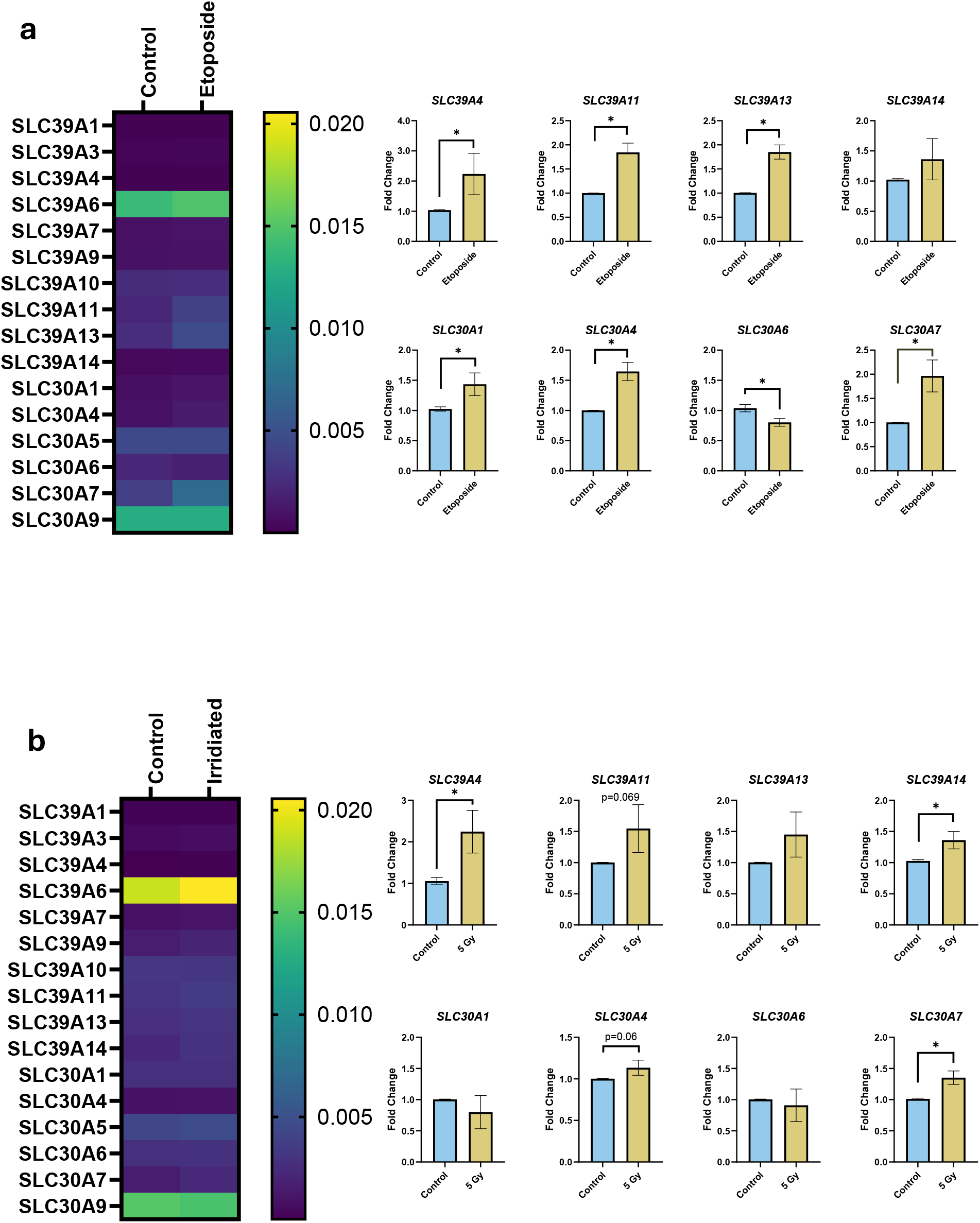
Zn transporter RNA expression in senescent C2BBe1 cells. RNA expression of key Zn transporters, measured by real-time quantitative PCR and normalized to *GAPDH* in etoposide-treated (A) or irradiated cells (B) at 48 hours of recovery. On the right are highly expressed or significantly changed genes in one or both senescence induction models. For analysis, a p < 0.05 was considered significant.

**Fig. 4.**
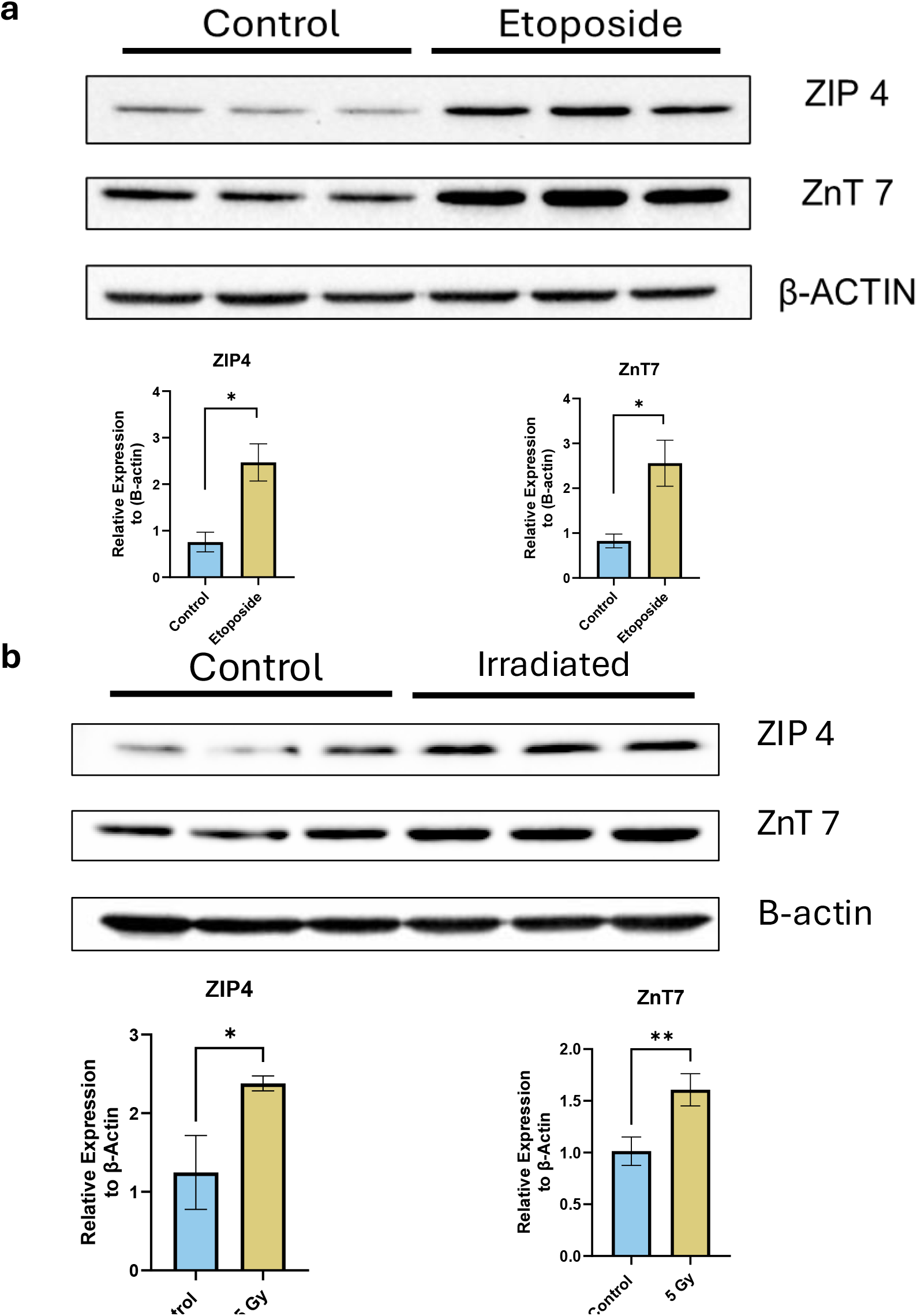
Western blot analysis of ZIP4 and ZnT7 protein expression in senescent C2BBe1 cells. Blots showing protein levels of ZIP4 and ZnT7 in whole-protein lysates from control and cells treated with etoposide (A) or irradiated (B). β-actin was used as a loading control. Protein quantification, measured in Imagelab, is shown below. For analysis, a p<0.05 was considered significant. Each band represents an individual biological replicate.

### Golgi Zn accumulation in senescent cells

While ZIP4 is well established to increase cytosolic Zn concentration in enterocytes [35], ZnT7 has been reported to transport Zn into the Golgi apparatus [40]. To determine whether ZnT7 upregulation leads to Zn accumulation specifically in the Golgi apparatus of senescent enterocytes, we utilized organelle-specific fluorescent probes and genetically encoded sensors. In the overlapped image of TR Ceramide, a Golgi-specific probe, and Zinpyr-1, we see that in both etoposide and irradiated cells, there was colocalization of the fluorescence, indicating Zn may be accumulating not only in the cytoplasm (Fig. 2), but also in the Golgi apparatus (Fig. 5A-B). To validate these findings, we employed a genetically encoded Zn sensor specifically targeting the Golgi apparatus. Baseline FRET ratio in etoposide-treated cells was higher, indicating increased Zn content in the Golgi apparatus, and when treated with Zn-pyrithione, there was less of an increase than that of the control cells. Upon the addition of a Zn chelator, both control and etoposide treated cell’s FRET ratio decreased. Interestingly, it seemed that the decrease was more rapid in the control cells (Fig. 6A). Due to the close relationship between the endoplasmic reticulum and the Golgi apparatus, we also examined Zn kinetics using an endoplasmic reticulum-specific sensor to test Zn levels. However, this did not show differences between the two groups (Fig. 6B) indicating that there are minimal alterations in the Zn-kinetics in the endoplasmic reticulum in senescent C2BBe1 cells. These results suggest that senescent enterocytes tend to accumulate Zn mainly in the Golgi.

**Fig. 5.**
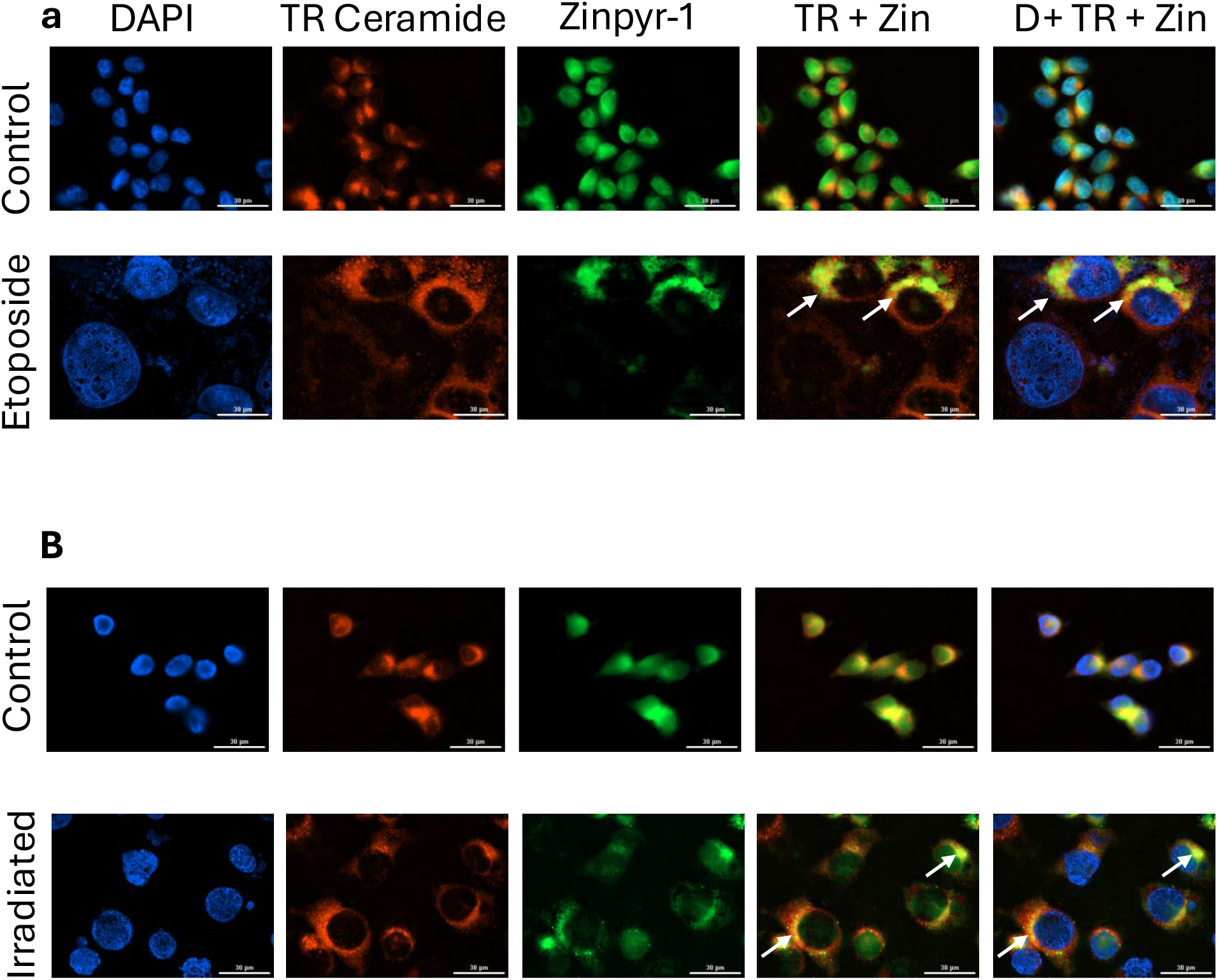
Fluorescent staining of Golgi apparatus and Zn in senescent cells. Representative images of control cells and etoposide-treated (A) or 5 gray irradiated (B) cells. Cells were tri-stained with BIPODY TR Ceramide, Zinpyr-1, and DAPI after 48 hours of recovery. Scale bar = 30 μm. Arrows indicate locations where Zn and Golgi apparatus markers overlap.

**Fig. 6.**
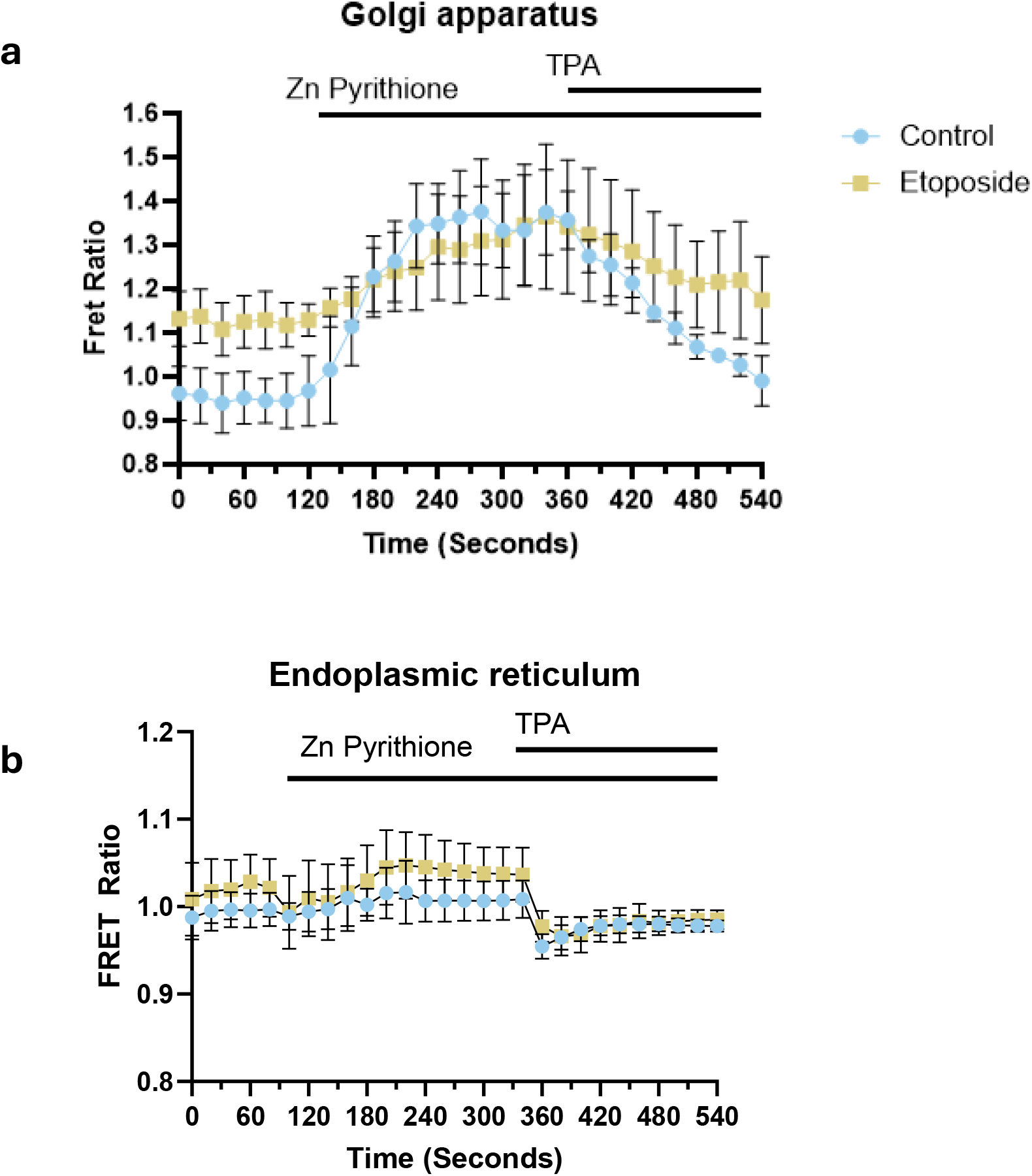
Zn kinetics in senescent C2BBe1 cells. Organelle-specific FRET sensors were transfected into cells, which were then treated with etoposide. Zn dynamics was measured using a LionheartTM FX fluorescent imager with administration of 20μM Zn pyrithione and 100μM Zn-specific chelator, TPA. Quantification of the FRET ratios for the Golgi apparatus (A) n=8, and endoplasmic reticulum (B) n=8 was performed using ImageJ software.

### Zn depletion during senescence induction decreases senescence-hallmarks

To examine the role of this Zn accumulation in senescence cells, we used a low dosage of TPA, a Zn chelator, in culture media during senescence development via etoposide treatment to prevent the Zn accumulation. To determine optimal TPA concentration, a viability assay was conducted, testing various TPA concentrations in both control and etoposide-treated cells. Above 0.15-0.175 µM TPA, control cells began to die off drastically (Fig. 7A). There was little effect on senescence cells’ viability, although above 0.5μM and higher seemed to cause complete loss of viability. Overall, the viability of senescent cells was not altered, likely due to anti-apoptosis mechanisms. Thus, we decided to use 0.15µM TPA, as it was the uppermost concentration that was still tolerated in the control wells. Upon Zn chelation, there was a rescue effect found in the SA-B-Gal staining (Fig. 7B). Examining the images, the morphology of the TPA and etoposide-treated cells was similar to that of the no TPA etoposide cells, which may indicate a similar morphology as typical senescent cells, but altered lysozyme activity. Looking at senescence-associated protein expression, there was a rescue effect in P21 expression; however, LMNB1 did not follow this trend (Fig. 7C). Gene expression for *CDKN1A, TIMP1, and SLC30A7* all experienced a rescue effect upon TPA treatment in etoposide-treated cells (Fig. 7D).

**Fig. 7.**
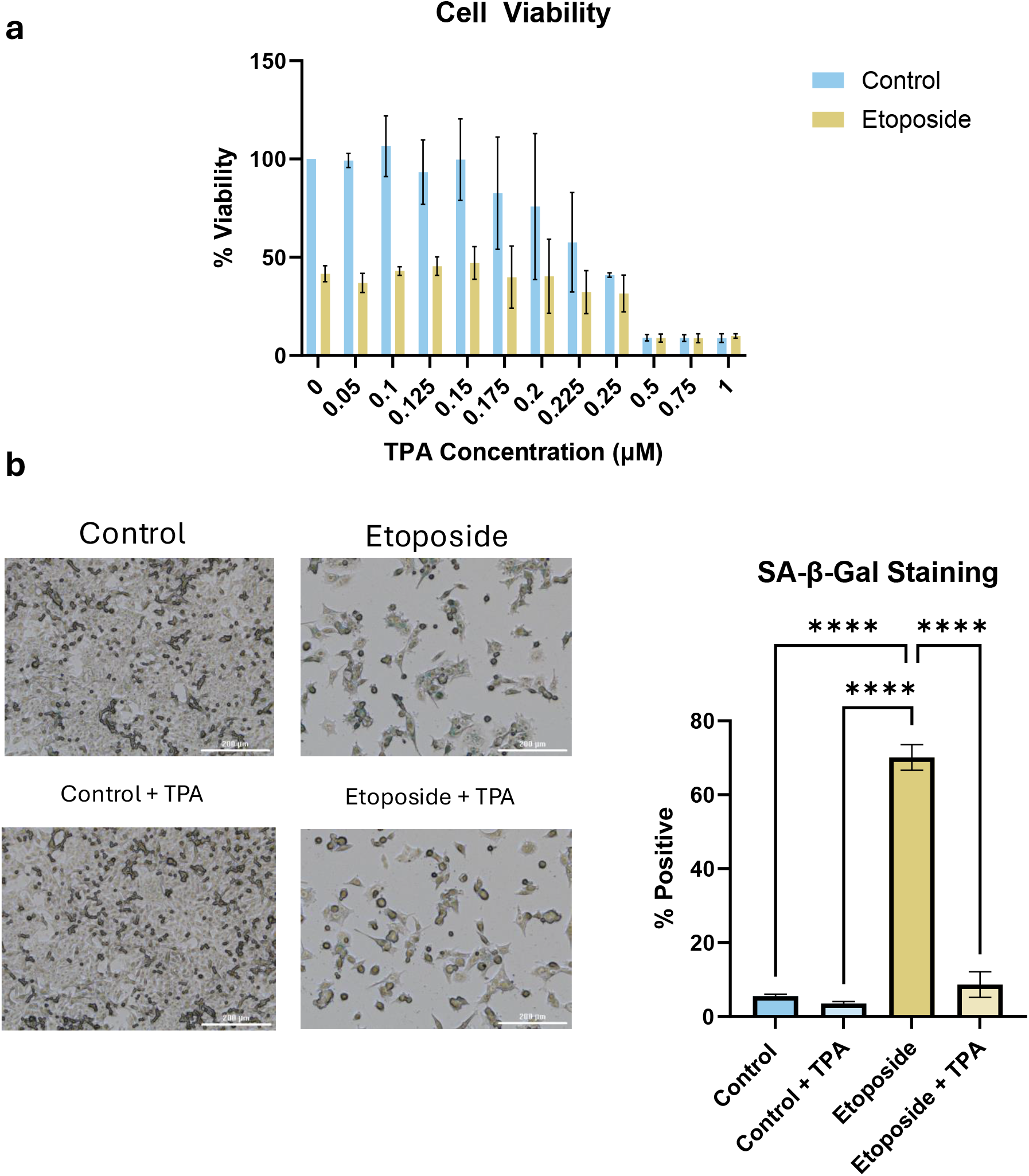
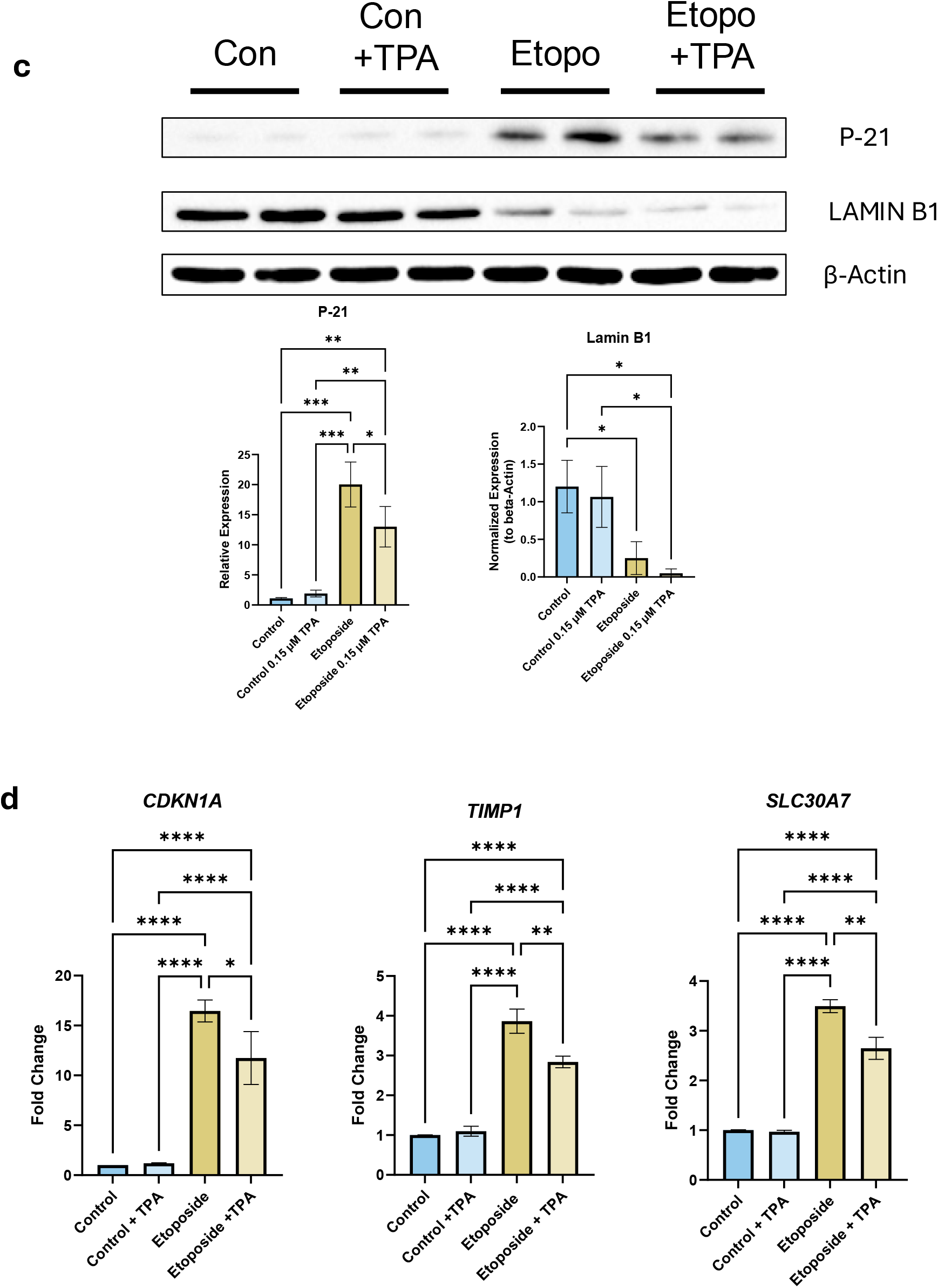
Zn chelation effects on senescent C2BBe1 cells. MTT cell viability assay in control and etoposide-treated cells treated with varying concentrations of TPA throughout the etoposide administration timeline (A). Representative images of senescence-associated ϐ-galactosidase staining at 48 hours recovery between control (top-left), control +TPA treated cells (bottom-left), etoposide (top-right), or etoposide + TPA (bottom-right), and their quantification (B). Scale bar = 200 μm. RNA expression *of CDKN1A, TIMP1*, and *SLC30A7*, measured by real-time quantitative PCR and normalized to *GAPDH* (C). Blots showing protein levels of p21 and LAMIN B1 in whole-protein lysates from control and cells treated with etoposide and/or TPA (D). β-actin was used as a loading control. Protein quantification was measured in Imagelab and is shown below. For all analyses, a p<0.05 was considered significant.

## Discussion

Various methods are used to induce senescence in cell models, including replicative senescence, DNA damage-induced senescence, oxidative stress, and media starvation. Regarding DNA damage-induced senescence, there are a plethora of methods involving direct DNA damage or DNA replication inhibitors; often, drugs or irradiation are used. We developed a method to induce senescence using etoposide by modifying a protocol from Kellers et al., which used Caco-2 cells [22]. Ionizing irradiation is another common method of inducing direct DNA damage in various cell types, and a dosage between 2-20 Gy is used depending on the cell line [29,30]. Common markers of senescence include the activation of p53 and/or p16INK4A pathways, morphological changes, lysosomal activity (SA-β-galactosidase), DNA damage response markers, mitochondria dysfunction/ROS production, loss of nuclear membrane structure (Lamin B1), and loss of proliferation [31]. In both of our models, multiple of these key hallmarks are seen, including p21 expression (indicating p53 pathway), loss of Lamin B1, increased SA-β-galactosidase activity, increased senescence-related gene expression, and loss of cellular proliferation. Data from this study support that there was senescence induction using both methods of induction. Using two methods of senescence induction allows us to highlight the effects of senescence, rather than treatment effects. Variation is seen across the modes of senescence induction; Thus, using the same cell line and two inducers of senescence, any matched outcome can be said to be due to senescence rather than the treatment. To this end, we see both methods induce similar responses in the markers of senescence. However, in Zn homeostatic genes, there is variation. By employing two methods, we could highlight our two Zn-related genes of interest.

Tightly regulated Zn concentrations are important for various cellular processes such as regulation of apoptosis, division, DNA replication, and protein and oxidative stress interactions. On an organismal level, Zn is critical for the immune system, growth and development, and neurological functions. Systemic Zn homeostasis is primarily maintained by regulating the amount absorbed at the small intestine. This study examined the relationship between Zn regulation and senescence, specifically in a cellular model for the intestinal epithelium.

Data from this current study are consistent with previous studies showing increased Zn accumulation with the development of senescence. Replicative culture-induced senescence in primary human fibroblasts (IMR-90s) and human umbilical vein endothelial cells (HUVECs) increases Zn concentration in the high passage cell groups compared to the young passages [19]. In human coronary artery endothelial cells (HCAECs), researchers observed disrupted Zn homeostasis upon replicative culture-induced senescence. Zn treatment in growth media (50uM) resulted in decreased cell viability and increased senescence markers. Furthermore, labile Zn increased over the culture lifespan in HCAECs and, in particular, senescent populations [32].

Functionally, restoring normal Zn concentrations to the aging organism, whether by dietary changes, improved absorption into the enterocyte, or improving transport through the epithelium layer into blood, will have impacts on the various systems Zn influences, such as improving memory and immune cell function [33,34]. Our data suggests that there is increased Zn accumulation in senescent intestinal epithelial cells, which may contribute to the progression of senescence and the decreased systemic Zn status seen with aging.

Zn transporters are essential for regulating cellular Zn homeostasis and have tissue, cell, and organelle-specific expression. ZIP4 is a highly expressed transporter in the GI system that is responsive to Zn availability and essential for maintaining Zn homeostasis.

Under Zn-deficient conditions, ZIP4 mRNA and protein levels increase to facilitate Zn uptake, a process mediated by the transcription factor Krüppel-like factor 4 (KLF4) [39]. Conversely, in Zn-replete conditions, ZIP4 mRNA expression is suppressed and the protein undergoes degradation to prevent excess Zn accumulation [36]. However, in our models, senescent cells displayed elevated ZIP4 expression despite high intracellular Zn levels, suggesting a disruption of the normal Zn-responsive regulatory mechanism. This aberrant ZIP4 upregulation was replicated in the enterocytes of aged mice and was rescued by gerotherapeutics such as rapamycin and metformin, which are known to reduce cellular senescence burden [37,38]. These findings imply that senescence may impair the Zn-dependent suppression of ZIP4, rather than falsely enhancing KLF4-mediated induction. As a result, the increased ZIP4 expression in senescent cells likely leads to Zn accumulation that cannot repress ZIP4 again and indicates a wider disruption of homeostatic gene regulation linked to cellular aging.

ZnT7 is localized in the pre-cis-Golgi apparatus, and functions to increase pre-*cis*-Golgi Zn, *trans*-golgi, and labile Zn in the cytosol [40]. Our data demonstrate increased ZnT7 expression in senescent cells, aligning with elevated Zn levels in the Golgi. Also, we observed that Zn chelation leads to a reduction in ZnT7 mRNA, suggesting that ZnT7 expression is positively regulated by cellular Zn levels. This suggests a possible feedback mechanism in which elevated Zn further increases ZnT7 expression, exacerbating Golgi Zn accumulation in senescent cells. Senescent cells exhibit structural alterations in the Golgi, including fragmentation and dispersion [41], as well as functional impairments such as altered glycosylation and vesicular secretion [42,43]. Given that Zn is an important cofactor of many Golgi resident glycosylation enzymes, the increased Zn status may contribute to Golgi dysfunction and impair proteostasis at the ER-Golgi interface [44]. This mechanistic link between Zn homeostasis and Golgi integrity in senescent enterocytes warrants further investigation.

While ZIP4 and ZnT7 were upregulated in both methods of senescence induction, there were other Zn transporters whose gene regulation upon senescence induction did not follow the same trends between etoposide and irradiation. HCAES and HUVECS with increased passages/replicative senescence experienced varied changes in their Zn transporter gene expression, with similar responses for *SLC30A1, SLC39A6, SLC39A13, SLC39A7*, and *SLC30A6*, with fold change for every gene besides SLC30A6 increasing with later passage [32]. Interestingly, *SLC30A7* significantly increased in late passage HUVECs, but decreased in HCAECs. As these cells are not closely related to C2BBe1 used in this study, the differences in lineages may explain the various impacts of senescence on specific gene expression.

Zn chelation in our study was performed using rather than classical high-affinity chelators such as TPEN or EDTA. TPA was selected due to its moderate affinity for Zn, which allows it to preferentially intercept biologically labile Zn pools without significantly disrupting tightly bound Zn in metalloproteins. Also, TPA has been reported to exhibit lower cytotoxicity compared to TPEN or EDTA, making it more suitable for long-term treatment during senescence induction. A low-dose regimen was used to minimize adverse effects on cell viability and proliferation, while preventing Zn accumulation during the development of senescence. Interestingly, we found decreased senescence markers, including SA-β-galactosidase, p21 protein and gene expression, as well as TIMP1, a colorectal cancer specific senescence marker [45]. However, cell viability and overall morphology were not rescued, suggesting that Zn chelation may selectively modulate certain senescence-associated pathways without fully reversing the senescent phenotype. For example, we did not find a statistically significant decrease in Lamin B1 expression in the TPA + Etoposide group.

Given the known role of Zn in regulating p21 expression and cell cycle arrest, our findings support a potential role of Zn in modulating senescence-associated signaling pathways. However, the incomplete rescue of senescence features highlights the limitations of global Zn chelation, particularly with agents like TPA that act across intracellular compartments. In this regards, whether Zn accumulation acts as a driver or a passenger in enterocyte senescence remains an open question, and future studies targeting organelle-specific Zn pools, such as those in the Golgi, may help clarify Zn’s mechanistic role in cellular aging.

This study sought to examine the role of senescence in gut epithelial cells and the impact it may have on Zn homeostasis. We found increased Zn levels in senescent cell populations, which may accumulate in the Golgi apparatus, and that Zn chelation during senescence development seems to reduce some senescence markers. Combining this data with *in vivo* models of aging Zn homeostasis, we can hypothesize that the senescent cells in the aging GI system may trap Zn and be an additional factor causing decreased Zn status in the aging organism.

## Abbreviations

ANOVA: Analysis of variance
HCAECs: Human coronary artery endothelial cells
HUVECs: Human umbilical vein endothelial cells
FRET: Flourescence Resonance Energy Transfer
TPA: tris(2-pyridylmethyl)amine
Zinc: Zn

## Statements and Declarations

The authors declare no conflicts of interest.

## Authors Contributions

The authors’ responsibilities were as follows: Kristofer Terrell and Sangyong Choi conceived the study; Kristofer Terrell wrote the manuscript with input and revisions from Sangyong Choi. Suyun Choi assisted with conceptual and methodological aspects of experimentation. Jiahn Choi assisted with the scRNA-sequencing data analysis. All authors read and approved the final manuscript.

## Acknowledgements

This work was partially supported by the Agriculture and Food Research Initiative (AFRI) of the USDA National Institute of Food and Agriculture (NIFA) (#2021-09056), the USDA Research Capacity Fund (#202425), and the Pilot and Exploratory Studies Core grant from the UConn Health Pepper Center Claude D. Pepper Older Americans Independence Center (5P30AG067988). We acknowledge the following laboratory members for their technical contributions: Ryan Lee, Alexis Bates, and Karina Huang. We sincerely thank the members from the University of Connecticut’s Environmental Health and Safety department, Peter Babin, Brianna Sullivan, and Amy Courchesne, for their assistance in utilizing the irradiator.

